# Single cell proteomics by mass spectrometry reveals deep epigenetic insight into the actions of an orphan histone deacetylase inhibitor

**DOI:** 10.1101/2024.01.05.574437

**Authors:** Benjamin C. Orsburn

## Abstract

Epigenetic programming has been shown to play a role in nearly every human system and disease where anyone has thought to look. However, the levels of heterogeneity at which epigenetic or epiproteomic modifications occur at single cell resolution across a population remains elusive. While recent advances in sequencing technology have allowed between 1 and 3 histone post-translational modifications to be analyzed in each single cell, over twenty separate chemical PTMs are known to exist, allowing thousands of possible combinations. Single cell proteomics by mass spectrometry (SCP) is an emerging technology in which hundreds or thousands of proteins can be directly quantified in typical human cells. As the proteins detected and quantified by SCP are heavily biased toward proteins of highest abundance, chromatin proteins are an attractive target for analysis. To this end, I applied SCP to the analysis of cancer cells treated with mocetinostat, a class specific histone deacetylase inhibitor. I find that 16 PTMs can be confidently identified and localized with high site specificity in single cells. In addition, the high abundance of histone proteins allows higher throughput methods to be utilized for SCP than previously described. While quantitative accuracy suffers when analyzing more than 700 cells per day, 9 histone proteins can be measured in single cells analyzed at even 3,500 cells per day, a throughput 10-fold greater than any previous report. In addition, the unbiased global approach utilized herein identifies a previously uncharacterized response to this drug through the S100-A8/S100-A9 protein complex partners. This response is observed in nearly every cell of the over 1,000 analyzed in this study, regardless of the relative throughput of the method utilized. While limitations exist in the methods described herein, current technologies can easily improve upon the results presented here to allow comprehensive analysis of histone PTMs to be performed in any mass spectrometry lab. All raw and processed data described in this study has been made publicly available through the ProteomeXchange/MASSIVE repository system as MSV000093434

**Abstract graphic:** 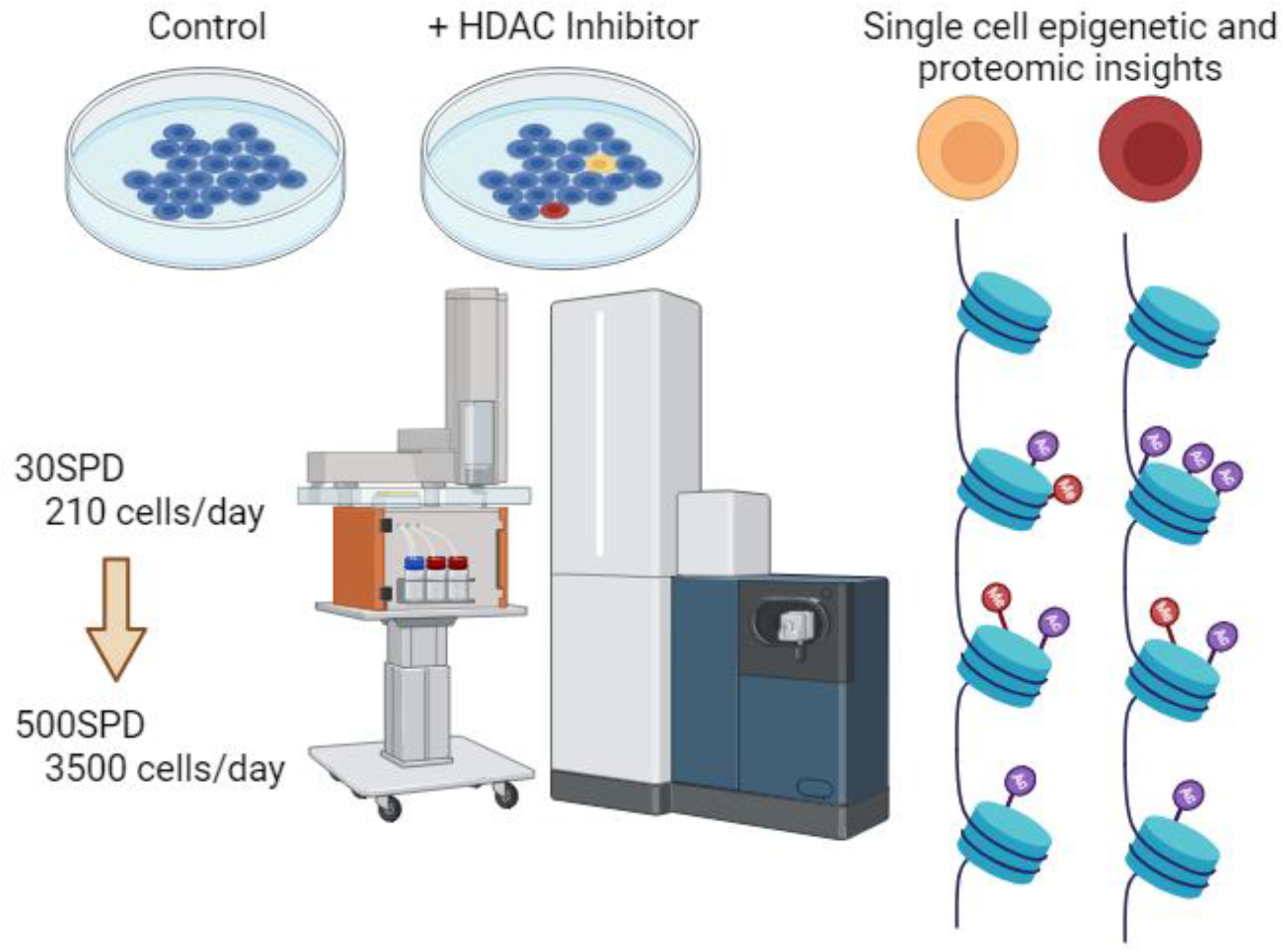

## Introduction

Histones are low molecular weight proteins that provide structural support for chromosomes and play well-characterized roles in gene expression. In addition, histones play roles in epigenetics, in part through crosstalk between their own protein post-translational modifications (PTMs) and chemical modification on the DNA itself.^1^ Acetylation of histones at key sites leads to open chromatin structure which permits transcription, while deacetylation causes a closed state and the corresponding repression of transcription. In addition, histone deacetylation has long been noted as a hallmark of cancer progression.^2^ As other characteristics of cancer progression are unchecked growth and cellular processes that occur despite damage that should stop these in normal cells, this relationship appears to be a linear one. A class of drugs loosely defined as Histone Deacetylase Inhibitors (HDACs) have shown promise in treating not only cancer, but also metabolic diseases, viral infections and age-related neurodegeneration. While promising, the core mechanisms and overall effects of many of these drugs remain poorly understood.^3^

Today, multiple methods exist which can profile cis-regulatory elements to help understand epigenetics in single cell populations. While these techniques are continuing to improve, they are largely derivates of ChIP-seq technology that use nucleotide, aptamer, or antibody probes specific for cis-regulatory elements. Newer variations of these methods can provide information on up to three histone PTMs as well as provide information on the nucleotide sequences with which they are associated.^4,5^ However, over 20 separate chemical PTMs have been identified on histone proteins by mass spectrometry that can exist in hundreds of possible combinations. Nearly all of these PTMs have been identified through the use of liquid chromatography mass spectrometry (LCMS).^1^

Work in our lab recently described the identification and quantification of 8 classes of protein PTMs in single human cells using multiplexed single cell mass spectrometry (SCP).^6^ In a follow-up work applied to understanding the KRAS^G12D^ inhibitor MRTX1133, histone PTMs were observed in nearly every one of more than 1,400 single human cancer cells analyzed.^7^ The main limitation in SCP by LCMS is the absolute protein concentration within each individual cell – more abundant proteins are exponentially easier to detect than lower abundance ones.^8^ As histone proteins exist in millions of individual copies per cell, peptides from all main classes of histones are easily identified in nearly every SCP study of nucleated cells.^9^ The signal quality from these proteins is so high that recent work described how the relative size of each single cell can be normalized solely by using the histone H4 protein abundance as a scaling factor.^10^ It seems likely, therefore, that drugs affecting histones, such as HDACs, could be studied by SCP. In addition, methods with higher relative throughput than those typically in use may still provide valuable insight into the effects of HDACs in single human cells, even when proteome coverage decreases as relative throughput increases. In this study, I test this hypothesis by analyzing pseudo-randomized control and HDAC treated single cells with multiplexed SCP methods allowing between 210 and 3,500 single cells to be analyzed per day.

## Results and Discussion

### An HPLC system designed for clinical samples can be applied to high throughput SCP experiments

The EvoSep One system is a fit for purpose LCMS system originally designed for clinical proteomics and other standardized high throughput proteomics applications.^11^ While limited to the validated methods provided by the vendor, the system provides value where reproducibility and throughput are a priority over method development. As described previously, our methods for multiplexed SCP allow for 7 cells to be analyzed per LCMS injection with the remaining three multiplex reagents used for blanks and controls.^7^ For clarity of presentation, I will refer to the methods in this study by the number of cells analyzed per day (CPD). For example, a 700 CPD method will allow, after blanks and carrier controls, 700 single human cells to be analyzed on one instrument in 24 hours. A summary of the results of analyzing single cells with various CPD methods is provided as **Table 1**. Surprisingly, doubling the throughput from 210CPD to 420CPD led to an overall increase in the average number of proteins identified in the study. As described previously, a commercially available labeled peptide digest was used for assessing the quantitative value of each method.^6,12,13^ The 210CPD and 420SPD methods resulted in nearly identical levels of observed isolation interference. While the 700SPD resulted in increased relative coisolation interference compared to the other two, the results for the met6 protein were still found to be quantitatively significant (**Supplemental Figure 1**). The 1400CPD, 2100CPD and 3500CPD methods resulted in less than 150 protein identifications in total and coisolation interference at levels which implied that any single cell measurements obtained were likely of no quantitative value (data not shown).

**Table 1.** A summary of proteomic and histone specific proteomic coverage of cells analyzed at each relative number of cells per day.

| LCMS injections/day (SPD) | 30 SPD | 60 SPD | 100 SPD | 200-500 SPD |
| --- | --- | --- | --- | --- |
| Column length | 15 cm | 8 cm | 8 cm | 4 cm |
| Cells per day after controls | **210CPD** | **420CPD** | **700CPD** | **1400-3500CPD** |
| Total protein groups | 1395 | 1755 | 964 | 171 |
| Average # proteins/cell | 342.97 | 453.51 | 215.52 | NA |
| Maximum # proteins/cell | 858 | 811 | 387 | NA |
| Unique histones proteins | 14 | 16 | 15 | 9 |
| Total histone PSMs | 4284 | 8022 | 3355 | 163 |
| Average % coverage of histones | 47% | 44% | 41.70% | 22.11% |
| Cells analyzed | 280 | 567 | 378 | 84 |

### Increasing SCP acquisition rates has less detrimental effects on histone measurements

LCMS based proteomics has always been biased toward proteins of the highest relative concentrations within a complex mixture. As such, most advances in increasing proteomic depth center on limiting repeatedly measuring peptides from the highest abundance proteins.^14,15^ Reflective of this, the number and average percent sequence coverage of histone proteins are only slightly diminished when increasing the number of samples analyzed per day. Even at 700CPD, when the number of proteins per cell has decreased by 54.9% relative to the 210 SPD method, the average percent coverage of histone proteins detected only decreases from 47% to 41.7%. Impressively, this trend continues even when increasing the sample acquisition rate to 3500CPD. Nine histone proteins were still detected, however, at only 22.11% sequence coverage of the histone proteins identified. As previously mentioned, isolation interference in the QC samples suggests that quantitative evaluation of these highest acquisition rate data is likely of negligible value.

### Sixteen individual combinations of histone PTMs can be identified in single cells using this approach

While multiple technologies today can monitor histone PTMs in single cells, these are currently limited to a maximum of three per cell, while most can monitor only one PTM on a single histone protein per study.^5^ In the approach described here, sixteen separate histone peptides can be identified with high confidence in single H358 cancer cells. **Table 2** is a summary of these peptide site identifications. Due to the high degree of homology in H3.1 and H3.3, the protein the PTM originated from could not always be determined. It should be noted that the method applied here utilizes a fully untargeted method for identifying peptides that are fragmented, sequenced, and subsequently assigned to each single cell. Targeted approaches or data independent analysis methods which are less biased by stochastic sampling should be applicable to epigenetic analysis of single cells while likely resulting in fewer missing values.^16,17^

**Table 2.** A summary of the histone PTMs identified with high confidence and assigned to single human cells in this study.

| Histone PTMs measured in this study | % of cells observed at 420 CPD |
| --- | --- |
| H1.2/H1.4 K34 Methyl (K35) Methyl | N/A |
| H1.3 K118 Methyl | N/A |
| H2B K21 Acetyl | 28.6% (166/580) |
| H3.1/H3.3 K15 Acetyl | N/A |
| H3.1/H3.3 K19 Acetyl K24 Acetyl | 16.2% (94/580) |
| H3.1/H3.3 K24 Acetyl | N/A |
| H3.1 K28 Methyl | 3.1% (18/580) |
| H3.1/H3.3 K28 Dimethyl | 66.3% (384/580) |
| H3.1/H3.3 K38 Dimethyl | 1.9% (11/580) |
| H3.1/H3.3 K80 Methyl | N/A |
| H3.1/H3.3 K80 Dimethyl | N/A |
| H4 K13 Acetyl | 14.1% (82/580) |
| H4 K13 Acetyl K19 Acetyl | 19.5% (113/580) |
| H4 K9 Acetyl K13 Acetyl K19 Acetyl | 10.0% (58/580) |
| H4 K80 Crotonyl | N/A |
| H4 T81 Phospho | N/A |

### Diagnostic ions are valuable complementary data in supporting histone acetylation sites

A valuable tool in the analysis of many PTMs is the presence of distinct fragment ions which correspond to modified amino acids or peptides. In glycoproteomic analysis the use of diagnostic oxonium ions is central to both instrument methods and to nearly all data analysis pipelines.^18^ As shown in **Figure 1** the well-characterized lysine diagnostic fragment ion of 126.09 can also be used as a secondary metric for the relative number of acetylation sites within histone peptide.^19^ A peptide identified as possessing a single acetylation site appears to produce a diagnostic ion roughly proportional to the reporter ion tag from a single cell (**Figure 1A**). As the number of identified acetylation sites in the peptide increase to two (**Figure 1B**) or three distinct sites (**Figure 1C**) the relative intensity of the 126.09 diagnostic ion scales in a manner roughly proportional to the reporter ion signal from each cell. While not truly quantitative, these ions provide supporting evidence for the identities of the peptides assigned by the search engine and mirrors recently described methods for analysis of intact glycopeptides.^20,21^

**Figure 1.**
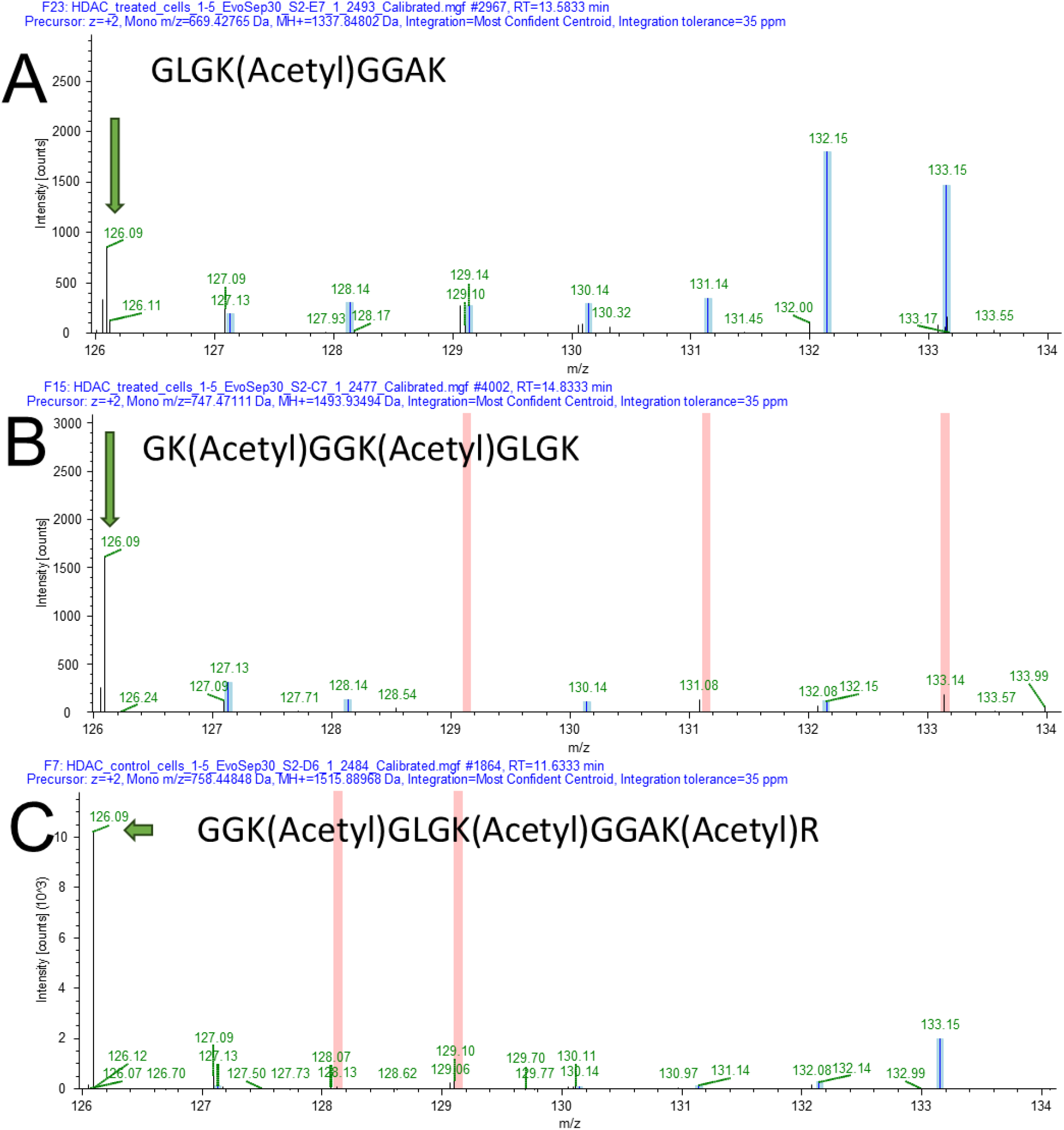
Lysine + acetylation diagnostic fragment ion intensity is roughly proportional to the number of acetylations observed within a peptide. **A**. A single acetylation site produces a diagnostic fragment ion (m/z = 126.09) with an intensity roughly equivalent to single cell reporter ion signal. **B**. A peptide with two annotated acetylation sites observed produces a 126.09 diagnostic ion with higher relative signal than any single cell reporter ions. **C**. A histone peptide with three acetylation further scales in relative intensity of diagnostic ion to single cell reporter ion signal. Blue highlights indicate where a reporter ion was successfully integrated to generate a quantitative value for the peptide within a single cell. Pink highlights indicate where no reporter ion was integrated.

For a more quantitative analysis of the use of the 126.09 diagnostic ion, I utilized the recently described DIDAR software to count the number of spectra with this ion from spectra identified as acetylated versus all spectra acquired. DIDAR reads LCMS files with MS/MS spectra and counts the number of spectra possessing user defined fragment ions.^22^ In the processed results from the 420 CPD method, 1,523 PSMs were identified as acetylated by the search software. These spectra were exported and were analyzed by DIDAR. DIDAR identified 349 processed fragmentation spectra (22.9%) possessing a fragment ion of 126.013 ± 0.005 Da. A strikingly similar 22.7% of spectra in the 210 CPD analyzed cells was likewise observed (175/769). Conversely, of 283,609 MS/MS original spectra obtained in total in the 420 CPD study only 1,687 spectra (0.59%) were found to contain this same ion under these same criteria. Similar numbers were found for the cells analyzed using the 210 CPD study. It should be noted that spotchecking the acetylated peptides manually as shown in **Figure 1**, suggests that the number of acetylated peptides with this diagnostic ion may actually be higher than the approximately 22% calculated by DIDAR. In order to process spectra from a TIMSTOF in Proteome Discoverer, we employ a previously described bin filtering system to reduce the overall noise in each spectra as one of the first steps in the analysis.^23^ In this case, only the top 12 most intense ions from each 100 Th window of each MS/MS spectra is retained for processing with Sequest. Due to the presence of the reporter ions in the first bin from 100 – 200 m/z, it appears likely that the 126.09 reporter ion is often filtered out. In our pipeline the reporter ion region is extracted from each spectra and analyzed separately with the quantification and identification values linked back together following data analysis. As the 126.09 reporter falls within the normal reporter ion region, this reporter ion region observed in post processing is a more accurate representation of the original mass spectra acquired. Despite this complication, it appears clear that spectra identified by as acetylated by our workflow are substantially enriched in the 126.09 acetylation diagnostic ion.

### Mocetinostat treated single cells have significantly higher levels of histone acetylation

Work in our lab has recently reported, and confirmed in a larger study, that post-translational modifications on high abundance proteins can be readily detected in many, if not all, single cells.^7^ To determine if PTMs could be meaningfully quantified following drug treatment NCI-H-358 cells (H358) were treated for 24 hours with mocetinostat, a class specific histone deacetylase inhibitor.^3^ A summary of some observations made using the 210 CPD method are shown in **Figure 2**. While no alteration in Histone H3 protein abundance was observed between the control and treated population (**Figure 2A**), two separate peptides annotated to possess acetylations were significantly increased in abundance. Both acetylation on K24 (**Figure 2B**) and a double acetylation on both K19 and K24 (**Figure 2C**) were found to be significantly increased following treatment. Similarly, four peptides were observed with a combination of acetylations affecting Histone H4 on K9, K13 and K17 and are, as a group, found to be highly significantly increased following drug treatment. However, I do observe a somewhat significant increase in total H4 protein abundance following mocetinostat treatment (**Figure 2D-E**). A recent study described the use of H4 protein levels as a proxy for normalizing between single cells of varying sizes.^10^ To test the effect that H4 protein abundance reflects on the increased acetylation observed, I reprocessed these data using the H4 scaling method. The increase in H4 acetylation remained highly significant (p<0.0001) following scaling (**Figure 3F**).

**Figure 2.**
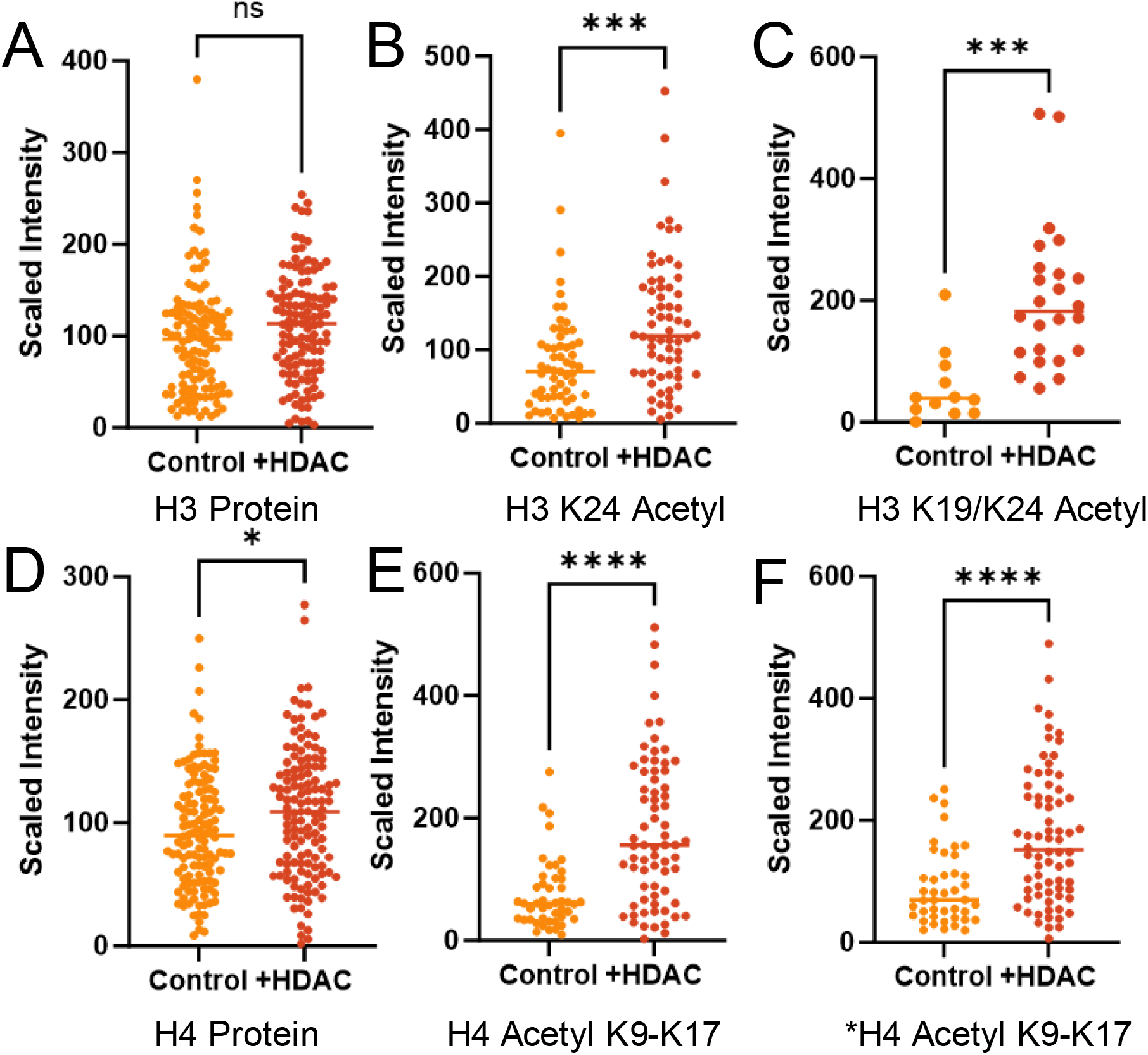
Mocetinostat treatment significantly increases the signal of acetylated peptides from histones H3 and H4. **A**. The measured abundance of the Histone H3 protein, as a whole, demonstrates no alteration following treatment. **B**. The H3 K24 peptide with acetylation as well as a double acetylation **C**. of K19 and K24 both demonstrate a highly significant increase in abundance following drug treatment. **D**. The Histone H4 protein, as a whole, does have a significant increase in total abundance (p = 0.0173) following treatment. **E**. A combined abundance plot of peptides demonstrating a combination of acetylation sites on K9, K13 and K17 are significantly increased following drug treatment. **F**. A reanalysis of E, after performing cell size normalization using H4 abundance. Source data are provided.

**Figure 3.**
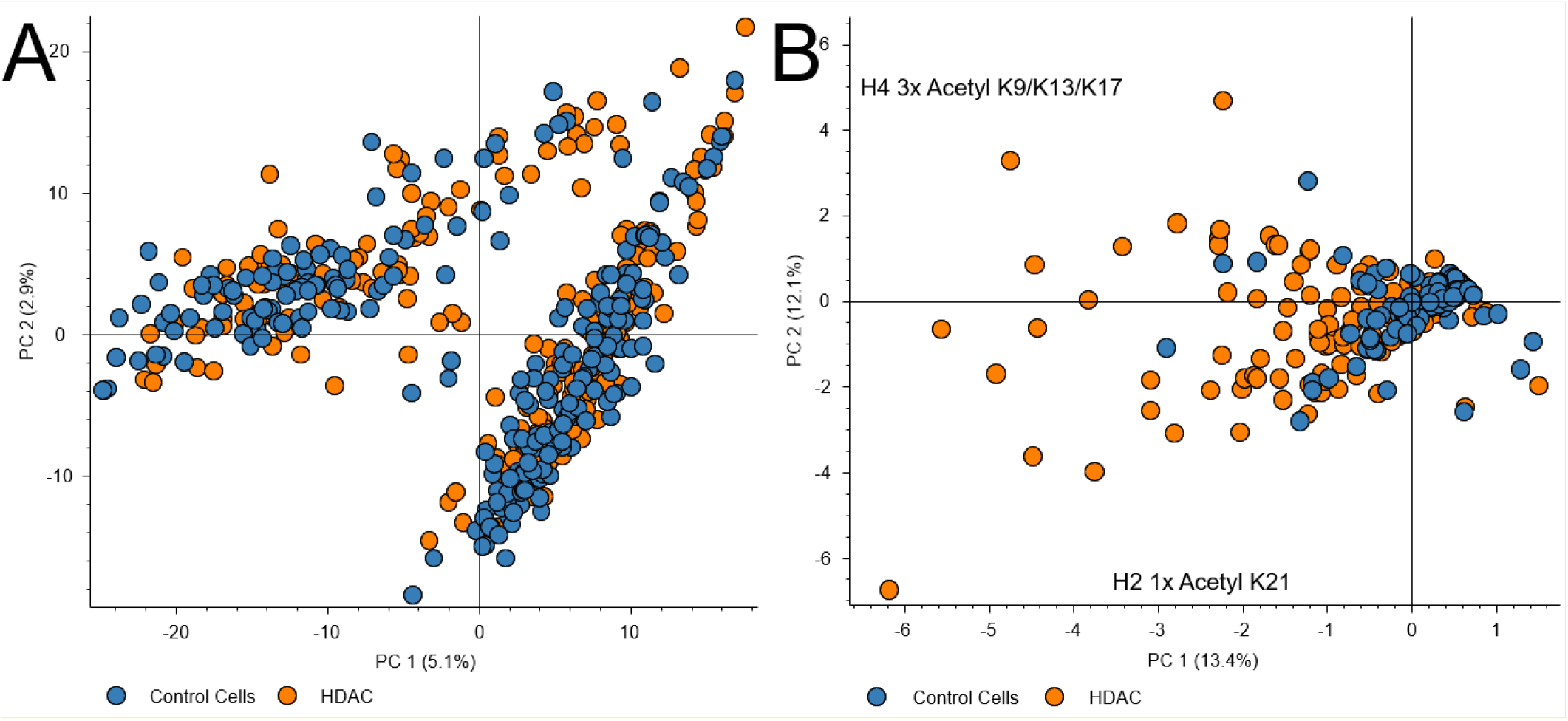
Single cell heterogeneity can be visualized by histone peptide signatures. **A**. A PCA plot of the protein abundance values for 580 peptides demonstrates only clustering by analysis batch. **B**. A PCA analysis of histone peptides from these same 580 cells with the large contributor to each dimension labeled.

Following the preprint of the first version of this manuscript, a comprehensive high depth proteomic analysis of 21 lysine deacetylase inhibitors was published by Chang *et al*., including an analysis of 10 concentrations of mocetinostat for 6 hours.^24^ To compare these results I reprocessed the 48 original LCMS files from the study using the same workflow used for the single cells in this manuscript, with minor alterations to account for mass analyzer architecture. In the bulk data, 52 unique histone acetylation sites were confidently identified, including every histone acetylation site reported in single cells in this study. A summary of the histone acetylation sites identified in bulk cell lysate as well as the relative abundance of each PTM at each dose of drug has been provided as **Supplemental Table 1**.

Additionally Chang *et al*., lends further support to the use of the 126.09 acetylation diagnostic ion as a confidence filter in the identification of acetylated peptides. In the reprocessing of these data for cells treated with mocetinostat, 80.7% (42/52) of histone peptides identified as acetylated possessed a clear 126.09 fragment ion in the diagnostic region.^24^ This is pertinent as the Orbitrap MS3 based method utilized in this study does not require prefiltering prior to analysis in this software and is likely more reflective of both the true prevalence of this diagnostic ion in unfiltered tandem mass spectra.

### Heterogeneity in histone PTM abundance can be observed following treatment

As previously noted, the proteomic alterations imparted by some drug treatments can not be readily observed in single cell proteomic data unless the alterations are relatively extreme. In the case of drugs where cell cycle alterations occur, such as treating KRASG12C mutant cells with a covalent inhibitor, a simple principal component analysis can readily separate control from treated cells.^25^ However, when using a noncovalent KRASG12D inhibitor at nonlethal doses, neither PCA nor T-SNE analysis alone could stratify control and treated cells.^7^ At the dose and time of mocetinostat treatment this is also the case when evaluating the cells in this study from the level of whole protein abundance measurements. The only clustering readily apparent in these data are the separation of LCMS batches analyzed approximately 1 week apart (**Figure 3A**). However, when performing a PCA analysis on the abundance of histone peptides, some stratification is apparent (**Figure 3B**). The loading plots for these principal components indicated that the largest factors contributing to these separations are both histone acetylation sites with Histone 2 K21 acetylation playing the largest role in PC1 and a tri-acetylation of histone 4 at K9, K13 and K17 playing the largest role in PC2.

### Proteomic data is obtained along with measurements of histone modifications in each cell

The use of standard proteomics statistical techniques is typically discouraged in single cell proteomics due to the inability of bulk protein statistics to accurately capture heterogeneity.^26^ It was therefore surprising to observe high significance in alteration in a single protein such as seen (**Figure 4A**) for S100-A9 (P06702). This calcium and zinc binding protein has, to the best of my knowledge, not been previously implicated in mocetinostat or other HDAC response mechanisms. Aside from keratins and proteins observed in a relatively small number of cells, the only other significantly altered protein was S100-A8. The proteins S100-A8 and S100-A9 are known to function as a hetero-tetramer complex in multiple contexts.^27^ As shown in **Figure 4B**, S100-A9 was observed as significantly increased in nearly every single cell studied at every acquisition speed. Surprisingly, the previously noted bulk proteomic study from Chang *et al*., on mocetinostat treated HeLa cells found no alterations in S100-A9 or S100-A8 expression at any dose of drug. This is curious as the human protein atlas lists the detection of the transcripts in neither HeLa or H358 cell line. However, both S100-A8 and S100-A9 have been previously linked to induced KRAS mutations in cell line experiments.^27^

**Figure 4.**
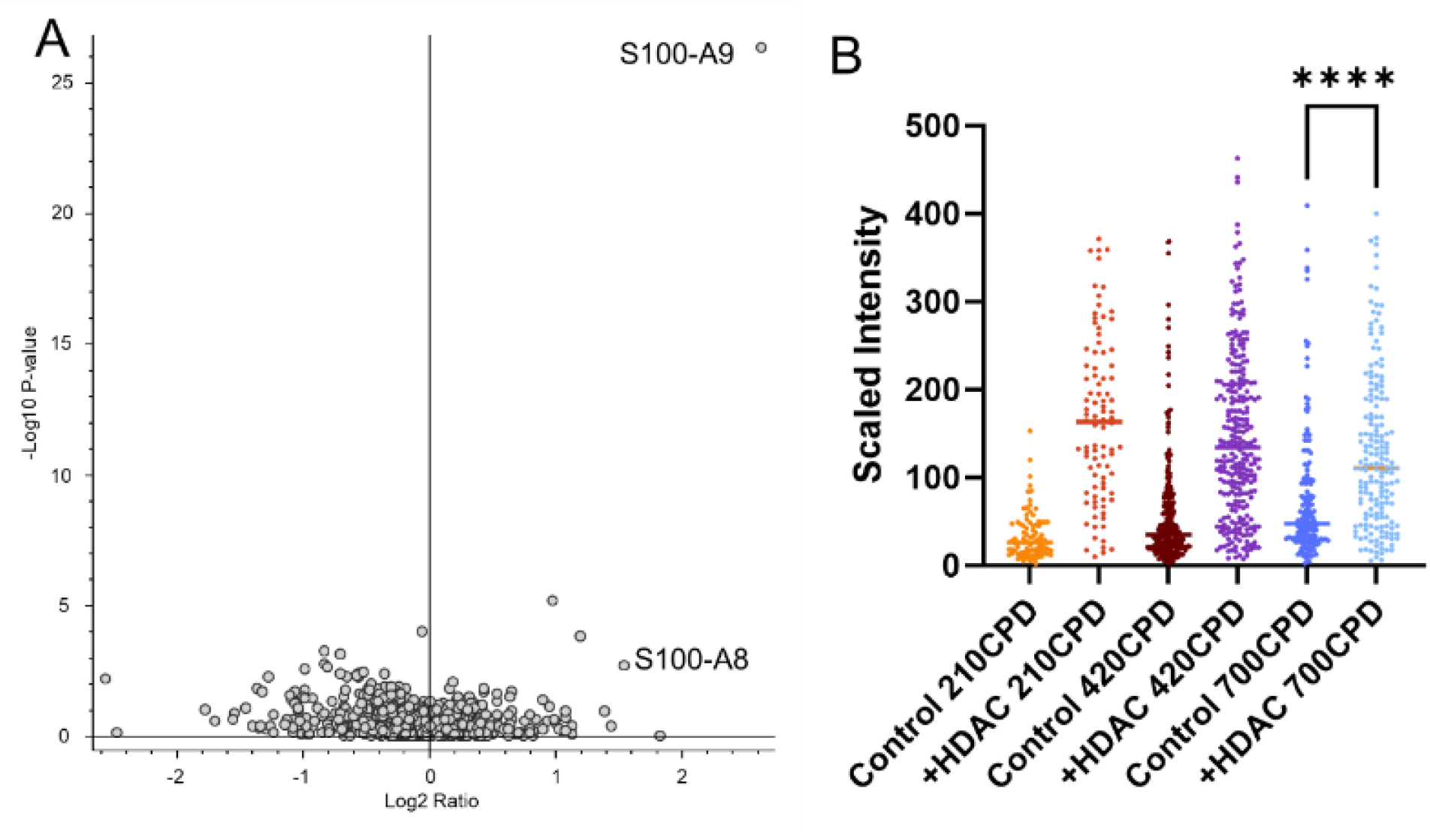
Mocetinostat treatment significantly increases S100A9/A8 protein in nearly all treated cells. **A**. A volcano plot from the 210 CPD analysis of control and treated H358 cells. **B**. Intensity values observed for the S100-A9 protein in all cells in this study. In all cases, a student’s t-test found a p-value < 0.0001 for control vs. treated cells. For this protein, regardless of the acquisition rate for the cells analyzed. The 700 cell per day method is labeled according to software defaults for pairwise significance. Source data are provided.

## Conclusions

There really is no question that LCMS is the method of choice for studying histone PTMs in bulk samples. Nearly every known modification on these important proteins has been identified by clever use of enzymes or derivatization coupled enzymatic digestion of these proteins.^28^ In addition, the incredibly high intracellular abundance of these proteins means that unless clever techniques to exclude them from analysis are employed, they will nearly always be sequenced with high coverage in every SCP experiment.^29,30^ Rather than ignore this important class or proteins, single cell epigenetics by mass spectrometry can exploit these characteristics to advantage. Global unbiased shotgun proteomics by data dependent methods always result in a high relative number of missing values, which is the entire argument for targeted, semi-targeted and data independent analysis methods.^31^ By simply changing the method described here for any of these other common LCMS proteomics techniques, the data in this study should be reproducible, and likely improved in terms of data completeness. In addition, while the highest CPD methods in this study were found to possess too much interference from background interference, this may be almost entirely due to mismatching of chromatography and methods. The 4 cm analytical columns used on this clinical HPLC system are intended for high speed, data independent methods utilizing wide windows to complement narrow chromatographic peaks.^32^ The use of relatively slow data dependent methods necessary here are simply a mismatch of technologies. Multiple new methods have recently been described that allow both high speed chromatography with higher relative chromatographic resolution, through the application of multi-pump systems which largely rely on the preparation of the next sample while the current sample is being analyzed.^33,34^ The application of these chromatographic methods may allow accurate quantification of histones and histone PTMs in single cells at throughput greater than 700CPD. Finally, while this study was intended to be focused almost entirely on the histone PTMs themselves, by using the global and untargeted approach, a previously unidentified response of cells to mocetinostat treatment was revealed. Ultimately, SCP is still a field of study in relative infancy compared to nearly any other -omics technique today, but the results presented here suggest that single cell epigenetics may be a fruitful avenue of future investigation.

## Methods

### Cell culture

The NCI-H-358 cell line (H358) was obtained from ATCC (CRL-5807) and cultured according to vendor instructions in RPMI-1640 (ATCC 30-2001) with 10% fetal bovine serum (ATCC 30-2020) and 10 mg/mL Penn Strep antibiotic solution (ATCC 30-2300). Single control cells were treated with DMSO or with 5 µm Mocetinostat (SelecChem S1122) prepared in DMSO. Treated cells were cultured alongside control cells for 24 hours prior to rapid washing of cells with ice cold magnesium and calcium free PBS (Fisher). The adherent cells were briefly rinsed in 3 mL of 0.05% Trypsin plus EDTA solution (ATCC 30-2001). This solution was rapidly aspirated off and replaced with 3 mL of the same solution. The cells were examined by light field microscopy and incubated at 37°C with multiple examinations until the adherent cells had lifted off the plate surface. The active trypsin was then quenched by the addition of 7 mL of the original culture media. The 10 mL solution was transferred to sterile 15 mL Falcon tubes (Fisher) and centrifuged at 300 *x g* for 3 minutes to pellet the cells. The supernatant was gently aspirated off and the cells were resuspended in PBS solution without calcium or magnesium with 0.1% BSA (both, Fisher Scientific) at 1 million cells per mL as estimated by bright field microscopy. Cells for single cell aliquoting were gently dissociated from clumps by slowly pipetting a solution of approximately 1 million cells through a Falcon cell strainer (Fisher, 353420) and the cells were placed on wet ice and immediately transported to the JHU Public Health sorting core. Non-viable cells were labeled with a propidium iodide (PI) solution provided by the core facility and briefly vortexed prior to cell isolation and aliquoting. Isolated single cells were deposited directly onto microwell plates containing two microliters of LCMS grade acetonitrile per well. A carrier channel of 150 single cells from control or treated cells were aliquoted into the first well in each row. The second well in each row was used as a method blank control well in which 20 nanoliters of the sorting buffer were aliquoted without transferring a single cell. Aliquoted plates were immediately transferred to dry ice prior to -80°C storage.

### Single cell isolation and preparation

Plates containing single cells were removed from -80°C in small batches and were immediately unsealed and placed onto a 95°C heatblock for approximately 90 seconds to fully lyse the cells and to dry off the remaining acetonitrile. The plates were cooled to room temperature and the lysates were digested with a solution of 100mM TEAB 0.1% DDM and 2 ng/µL sequencing grade trypsin. Two µL were added to each blank and single cell well and 4 µL was added to each carrier channel well. The plates were briefly centrifuged to remove air bubbles and were tightly sealed with plate sealing tape. The digestion was allowed to proceed at room temperature (approximately 18°C) overnight. Following digestion, the plates were thoroughly centrifuged and were labeled with TMTPro reagent. For this experiment, the carrier channel was labeled with TMTPro 135, the method blank control was labeled with 126 and the single cells were labeled with 127c, 128c, 129c, 130c, 131c, 132c, 133c and 134c. Due to impurities from the carrier channel, the 134c channel was ignored in all downstream analyses. Control and treated plates were combined using a previously described method for pseudo-randomization.^7^ The result was that in each LCMS experiment, the carrier channel contained an equal mixture of peptides from control and treated single cells (approximately 75 cellular volumes of each) and each injection contained both control and treated cells. The loading method ensures that in each subsequent injection, the cells analyzed are reversed. For example, in injection 1, channels 127-130 would contain control cells, where 131-134 would be treated cells. In the next injection these would be reversed, in that 127-130 were treated cells and 131-134 control cells. Each injection was labeled in a manner to allow original cellular identities to be deconvoluted in downstream data analysis.

### LCMS instrument parameters

An EvoSep One (EvoSep) and TIMSTOF SCP (Bruker Daltronic) was used for all analyses. EvoTips were prepared identically for all experiments resulting in each tip containing approximately 30 nanograms of total labeled cells if making the ludicrous assumption that no sample loss occurred in sample preparation. The EvoSep one was operated using PepSep columns coupled to a 10 µm CaptiveSpray emitter through a “Zero Dead Volume” steel union. The TIMSTOF SCP was operated in ddaPASEF mode using a 75 millisecond ramp time, with 8 ramps per cycle. A target of 20,000 ions was used with a minimum threshold of 2500 counts. TIMS Stepping was employed in which the low mass fragment ions were obtained using a collision energy dependent on 1/k0 value ranging from 45-90. Peptide sequencing data was obtained using a CE of 21-60. A 1/k0 isolation window of 0.7-1.45 was used and a custom polygon was used to attempt to collect only +2 or greater ions with an m/z of 380 or higher.

### Data Analysis

As previously described, I utilize a single point recalibration method using the 135n carrier channel signal to adjust the reporter ion region using an in-house developed tool (pasefRiQ Calibrator). This secondary calibration allows tighter mass accuracy tolerances to be used during the final data analysis, resulting in reduced background noise.^6^ For all control and treated cells a linear mass shift of -0.00564 from 100-136 m/z. The recalibrated output files were processed in Proteome Discoverer 2.4SP1 using SequestHT and Percolator.^35^ Briefly, the MS/MS spectra were binned into 100 Da segments and filtered so that only the top 12 most abundant ions from each bin were retained. The resulting cleaned spectra were searched with a 15ppm MS1 tolerance and 0.03 Da MS/MS tolerance. TMTPro labels were considered static on the N-terminus and dynamic on lysines to allow for the search for lysine PTMs. Methionine oxidation was the only additional dynamic modification. The default cutoffs for Percolator PSM validation and peptide and protein FDR determination were employed in all analyses. Reporter ions were integrated using a 35ppm mass tolerance window and quantification values were only used for unique peptides. A sum based normalization of all PSM signal for TMTPro channels 127-133 as well as the raw non-normalized abundance values. For cell size normalization, the H4 protein was used as the sole scaling factor for a secondary analysis of the 420SPD data. For reanalysis of the Chang et al., bulk proteomics data, the same pipeline was utilized, with the exception that a 10 ppm and 0.6 Da MS1 and MS2 tolerance, respectively, were utilized and the TMT6-plex label was appended as the modifications. All 48 fractions offline fractions were analyzed as fractions for the sake of quantification and ratios were produced by comparing each dosage point to the mock treated cells.

## Data Availability Statement

All Bruker. d files and Proteome Discoverer processed data has been deposited at the MASSIVE.UCSD.EDU public repository and is publicly available as accession MSV000093434.

## Acknowledgements

I would like to thank Hao Zhang of the JHU Public Health sorting core for invaluable assistance in this and every other single cell proteomics study performed in my lab. In addition, a big thank you to Ahmed Warshanna and Tarsh Shah for help preparing these cells. I would also like to thank Cynthia Wohlberger, Amanda Smythers, and Ben Garcia for answering naïve questions about histones and their analysis.

## Author Contributions Statement

Conceptualization: BCO

Methodology: BCO

Investigation: BCO

Funding acquisition: BCO

Writing – original draft: BCO

Writing – review & editing: BCO

## Supplemental Materials

**Supplementary Figure 1.**
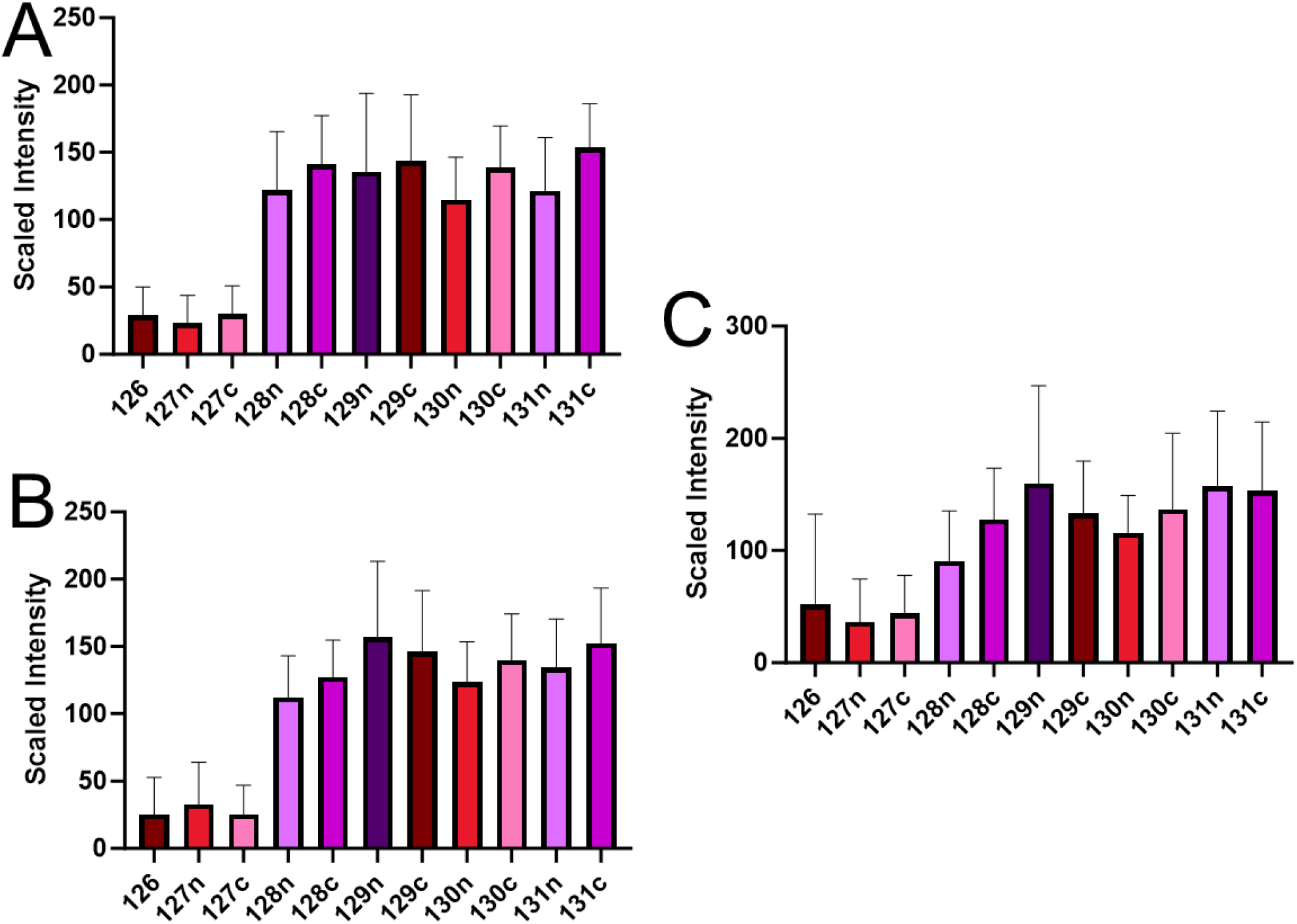
Summary of peptide abundance observed for 20 nanograms of TMT labeled yeast digest commercial triple knock out standard for Met6 where 126, 127n and 127c are Δmet6 knockout strains. **A**. A summary of peptides from the 210 CPD method. **B**. A summary of the 420 CPD method. **C**. Summary results from the 700 CPD method. Source data are provided.

